# Spatial exclusion leads to tug-of-war ecological dynamics between competing species within microchannels

**DOI:** 10.1101/2023.01.10.523527

**Authors:** Jeremy Rothschild, Tianyi Ma, Joshua N. Milstein, Anton Zilman

**Affiliations:** Department of Physics, University of Toronto, Ontario, Canada; Department of Chemical and Physical Sciences, University of Toronto Mississauga, Ontario, Canada; Institute for Biomedical Engineering, University of Toronto, Ontario, Canada

## Abstract

Competition is ubiquitous in microbial communities, shaping both their spatial and temporal structure and composition. Many classic minimal models, such as the Moran model, have been employed in ecology and evolutionary biology to understand the role of fixation and invasion in the maintenance of a population. Informed by recent experimental studies of cellular competition in confined spaces, we extend the Moran model to explicitly incorporate spatial exclusion through mechanical interactions among cells within a one-dimensional, open microchannel. The results of our spatial exclusion model differ significantly from those of its classical counterpart. The fixation/extinction probability of a species sharply depends on the species’ initial relative abundance, and the mean time to fixation is greatly accelerated, scaling logarithmically, rather than algebraically, with the system size. In non-neutral cases, spatial exclusion tends to attenuate the effects of fitness differences on the probability of fixation, and the fixation times increase as the relative fitness differences between species increase. Successful fixation by invasive species, whether through mutation or immigration, are also less probable on average than in the Moran model. Surprisingly, in the spatial exclusion model, successful fixations occur on average more rapidly in longer channels. The mean time to fixation heuristically arises from the boundary between populations performing either quasi-neutral diffusion, near a semi-stable fixed point, or quasi-deterministic avalanche dynamics away from the fixed point. These results, which can be tested in microfluidic monolayer devices, have implications for the maintenance of species diversity in dense bacterial ecosystems where spatial exclusion is central to the competition, such as in organized biofilms or intestinal crypts. The results may be broadly applied to any system displaying tug-of-war type dynamics with a region of quasi-neutral diffusion centered around regions of deterministic population collapse.

**Author summary:** Competition for territory between different species has far reaching consequences for the diversity and fate of bacterial communities. In this study, we theoretically and computationally study the competitive dynamics of two bacterial populations competing for space in confined environments. The model we develop extends classical models that have served as paradigms for understanding competitive dynamics but did not explicitly include spatial exclusion. We find that spatial effects drastically change the probability of one species successfully outcompeting the other and accelerates the mean time it takes for a species to exclude the other from the environment. In comparison to the predictions of population models that neglect spatial exclusion, species with higher selective advantages are less heavily favoured to outcompete their rival species. Moreover, spatial exclusion influences the success of an invasive species taking over a densely populated community. Compared to classical well-mixed models, there is a reduction in the effectiveness of an invaders fitness advantage at improving the chances of taking over the population. Our results show that spatial exclusion has rich and unexpected repercussions on species dominance and the long-time composition of populations. These must be considered when trying to understand complex bacterial ecosystems such as biofilms and intestinal flora.

## Introduction

Ecological competition is a ubiquitous feature of multi-species communities. It often manifests itself through direct antagonistic interactions between species, such as bacterial toxins, metabolic waste products and parasitic infections [1–3]. Competition also commonly occurs indirectly through various exploitative scenarios that deplete communal resources. Computational models of the dynamics of populations, framed in the context of a competition for finite system resources (e.g., light, food, population density, etc.) [4–12], have defined various heuristic measures of this competition for resources, such as the niche overlap, competition strength, and carrying capacity. Although these measures are commonly used to describe the dynamics of population growth and co-existence, a deeper understanding of the processes that govern the structure of ecological communities is acquired by exploring the mechanisms of the resource competition that underlie these coarse-grained, aggregate parameters [13–18].

Among the various resources required for population maintenance and growth, physical space is essential for expansion and access to additional resources [19–21]. In fact, individuals inherently require physical space for both their own growth and those of their progeny. Spatial competition can result in complicated patterning, synchrony of population distributions, spatial segregation into different niches within the environment and hosts, as well as other non-trivial dynamics [20, 22–25].

In bacterial communities, a variety of spatially ordered configurations may emerge from similarly distributed initial populations. This spatial structure plays an important role in medicine, industrial fabrication, and food production [26–30]. In bacterial biofilms, for instance, microbial populations form complex structures wherein various species segregate [31–33]. Different layers of bacterial species within a biofilm may have different sensitivity and resistance to antibiotics that restrict our ability to treat associated infections [34, 35]. The biogeography of bacteria in the digestive tract, which form the human digestive microbiome, illustrates another spatially heterogeneous ecology [36, 37]. In particular, the intestinal tract hosts diverse microbiota whose complex physical structures, such as mucus densities and epithelial crypts, have direct implications on the long-term composition of the bacterial community [38]. It is, therefore, necessary to understand how spatial constraints, arising from a confining environment and crowding/exclusion by other bacteria, shape the dynamics of each species and the overall patterning of the populations.

Spatial constraints may also have ramifications for the overall ecological diversity. For instance, the boundary between expanding fronts of different bacterial populations, grown on solid substrates, fluctuates superdiffusively. This encourages accelerated genetic drift that may limit diversity more rapidly than neutral mutation models void of any spatial dynamics [39]. Alternatively, diversity may be increased in ecosystems wherein species undertake differing strategies in relation to the space they occupy - sometimes referred as distinct spatial ‘niches’. For instance, trade-offs between motility and competitive ability may allow for coexistence between competing species [40].

It is well known that diversity may be strongly influenced by the invasion of external species. As initially noted by Gause *et al*., extinctions are frequently observed in a closed competitive ecosystem within a laboratory setting even though the similar ecosystem persists indefinitely in nature [41–44]. This suggests that invasion events, which implicitly rely on a partitioning of the space between the local and meta-community, contribute crucially to the population dynamics by reintroducing individuals into the ecosystem [45– 48]. For instance, persistent diversity is observed through the fragmentation of continuous ecosystems in studies of patch-models of ecology and theories of island biogeography [16, 49–52].

The contest for space is critical in environments with small total populations, which accentuates the individual composition of the colony, such as within intestinal crypts [53, 54]. In recent years, microfluidic devices have started to provide controlled systems for exploring the population dynamics of small bacterial microcolonies. Experiments in microfluidic monolayer devices, of various geometries, have shown that small populations of asymmetric bacteria, like *E. coli*, can align into highly ordered arrangements [55]. These populations behave differently from their well-mixed counterparts given the densely packed nature of their confinement [56, 57]. Cell morphology and the confining geometry are observed to greatly affect the ordering and fixation probability of cell populations in these devices [57–62].

Certain models, like the classical Moran model, subsume spatial factors into their framework by assuming that the species are well-mixed (i.e., that species abundances are uniformly distributed in a certain location). Similarly, other models of bacterial competition that incorporate space more explicitly also utilize the well-mixed assumption to explain ecological processes, such as the spatiotemporal synchrony of densities of different species [63–65]. However, well-mixed continuous models that describe the concentration of bacterial populations generally neglect explicit spatial exclusion. Although many factors may influence competition for space, these densely packed populations must fundamentally exclude each other through mechanical interactions between cells. Consequently, modeling the competition of bacteria in small, confined environments requires explicit consideration of the system’s spatial configuration to correctly describe the competitive dynamics.

In this paper, we explore how spatial constraints imposed by cellular interactions influence the population dynamics of competing microcolonies of bacteria. Inspired by the experimental setting of a microfluidic chemostat, we investigate how two species of bacteria compete through physical exclusion in an open, single lane microchannel [66]. We first review a general model of competing populations that reduces to the Moran model for well-mixed populations, before describing an extension to this model that incorporates spatial exclusion. We calculate the probabilities and the mean first-passage times of fixation for the spatial exclusion model, comparing the results to the Moran model. We then explore how fitness differences between the two species - for instance, differences in the bacterial growth rate or doubling time - shape the competitive landscape by affecting the time to and probability of fixation. Finally, the model is used to investigate invasion events that perturb a local community and the characteristics of a successful invasion.

## Models and Methods

We characterize the state of the system by the abundances of the competing species (i.e. the numbers of individuals of each species found in the ecosystem) which reflect the dynamics and the evolution of the community. We focus on the processes where the competition amounts to a zero-sum game: different populations compete for dominance under a constraint of finite total population size, *N*, determined by limitations of the inhabited space. Thus, in a two species system with a finite total population size studied here, fixing the abundance of the population of the first species, *n*, determines the abundance of the population of the second species, *N* − *n*, and both species compete to maintain their non-zero abundances in the system. The constraint of a fixed population size means that the dynamics of this 2 species system can be mapped onto a one-dimensional process defined by the abundance *n*.

Several important models have been used to study the effects of different competition mechanisms on the species abundance and the community structures. A classical, highly influential model - the Moran model (and its variants) - has served as the paradigm for understanding the effects of stochastic ecological drift and natural selection on the diversity of a well-mixed population [59, 67–70]. A closely related model, Hubbell’s neutral theory of biodiversity has been used to describe the emergence of the species abundance distribution in a neutral immigration-birth-death process [71]. Among others, Lotka-Voltera models further explore the role of species interactions and niche overlap on the interspecific competition; their frameworks can also be roughly mapped to the Moran model in their neutral regimes [16, 72, 73]. The fundamental stochastic nature of the ecological processes underlies all these models, where stochastic fluctuations of the abundances emerge from the demographic noise (i.e., the inherent randomness of birth and death events in a population).

We investigate the population dynamics of two competing species as a discrete stochastic process denoting the probability of being in a state with one species abundance at *n* (and the other species abundance at *N* − *n*) at time *t* is *p*(*n, t*). The population abundance of a species can change either through births or deaths of the individual cells, with the probability of a birth or death in the population *n* in an interval of time Δ*t* denoted as *T* ^+^(*n* → *n* + 1, Δ*t*) or *T* ^−^(*n* → *n* − 1, Δ*t*), respectively [74, 75]. The evolution of the probability *p*(*n, t*), is governed by a one-dimensional forward master equation (ME)

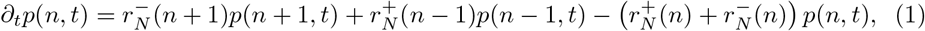

where 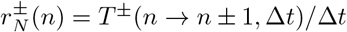 are the transition rates for events of an increase or decrease in abundance [76].

We are interested in the process of fixation wherein the abundance of one species approaches *N*, effectively outcompeting the other species by removing it from the system. This fixation can be viewed as a first-passage process that occurs when the abundance of a species reaches either of the absorbing states, at *n* = 0 and *n* = *N*, at which point the system settles at steady-state with one species dominating indefinitely [16, 77]. In these processes, the first-passage probability and the mean first-passage time (MFPT) are characteristics of the system which elucidate the dynamics of the process [75].

The mean-first passage time to either fixation state, from a starting abundance *n, τ* (*n*), relates the average time the competition between the two species lasts before one takes over and is described by the backward equation

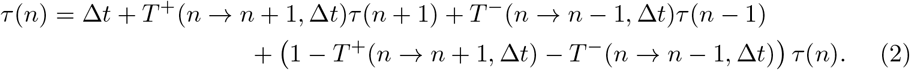

The Fokker-Planck (FP) expansion to order 𝒪(*N* ^−2^) of this equation is

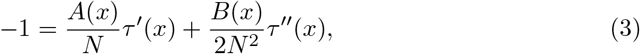

where *A*(*x*) = *r*^+^(*x*) − *r*^−^(*x*) and *B*(*x*) = *r*^+^(*x*) + *r*^−^(*x*) with the transformation 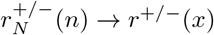 and *x* = *n/N*. A similar FP expansion in 1*/N* of the first-passage probability to settle in either absorbing abundance state *f* ∈ {0, 1} results in a continuous description (see S1 Appendix)

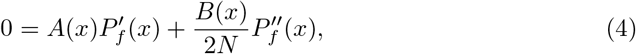

where *P*_*f*_ (*x*) is the probability of being absorbed at the state *f* starting from an abundance *x*. For instance, *P*_1_(*x*) is the probability that a species with relative abundance *x* will fill the space to fixate at the absorbing abundance *x* = 1.

Finally, the corresponding equation for the directional MFPT of fixation through one of the absorbing states *f* ∈ {0, 1} may be written as follows:

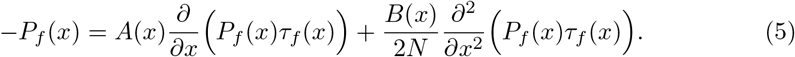

The solutions to these equations depend on the choice of the birth and death rates of the model. In the classical Moran model, which represents mixed populations without spatial structure, a random individual from a fixed and finite population of size *N* is selected to give birth at each time step while, simultaneously, a random individual is selected to be removed from the system to make room for the progeny and maintain a constant total population *N* (see Fig 1A). In the truly neutral case without fitness differences between the species, the probability for the species with *n* individuals to increase in size is the product of the probability that a member of that species is selected for a birth event (*n/N*) and the probability that the other species is selected for a death event ((*N − n*)*/N*). Consequently, the probabilities of abundance transitions in a discrete timestep Δ*t* in the Moran model are *T* ^+^(*n* → *n* + 1) = (*n/N*)((*N* − *n*)*/N*) and *T* ^−^(*n* → *n* − 1) = ((*N* − *n*)*/N*)(*n/N*)). These discrete time probabilities are transformed and rescaled to represent the rates used in Eq 1, 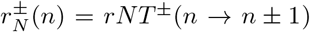 where *r* = 1*/*(*N* Δ*t*). Plugging these expressions into Eq 3 and Eq 4 in the continuum limit *x* = *n/N* recovers the classical Moran model result, namely, the probability of a successful fixation is equal to the relative initial abundance of the fixated species, *P* (*x*) = *x*, and the mean time to fixation is *τ* (*x*) = − *N/r*[(1 − *x*) log(1 − *x*) + *x* log *x*] [78].

**Fig 1.**
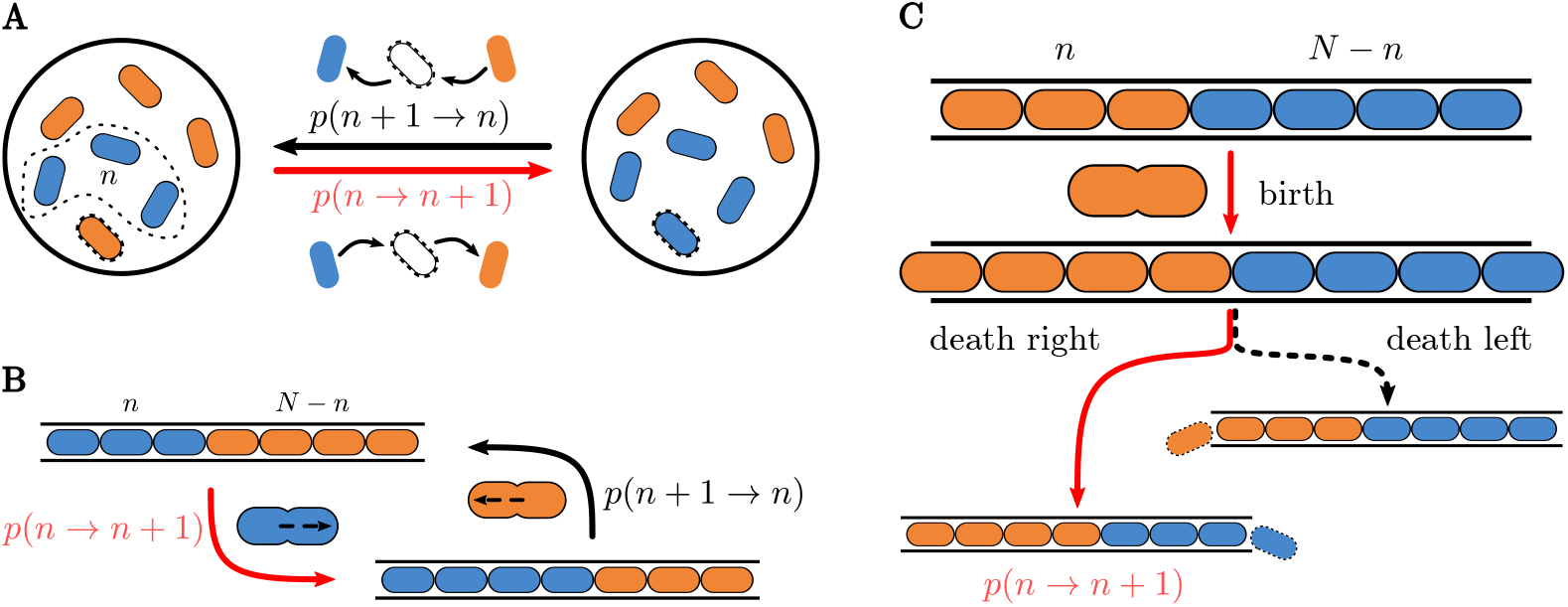
Illustration of minimal models of ecological competition. **(A)** Illustration of the classical Moran model within a well-mixed population of two species of bacteria. Change in the abundance, *n*, of a species relies on selecting an individual of that species to give birth while simultaneously having an individual of the other species die. **(B)** A spatial exclusion model for 1-dimensional competition within a micro-channel. Instead of a well mixed population, the cell populations are segregated, each to one side of the open channel. **(C)** Within the spatial model, a birth by one species may be followed by a death by either species, resulting in either a state transition with probability *p*(*n* → *n* +1) or no change at all. The probability of dividing to the right or left is weighted by the location of the dividing cell in the channel.

Fitness differences/selective advantages between the species can be incorporated into the model by appropriately changing the probability of selecting a species for a birth/death. By assuming that a fitness difference *s* - or coefficient of selection in evolutionary dynamics - exists between the species, the rates of the Moran model with selection are

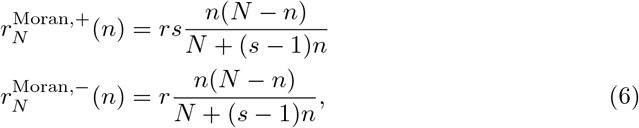

where *s >* 1 indicates a fitness advantage. However, neither the neutral nor the selective Moran model account for the spatial consideration of physical exclusion.

Building on this paradigmatic model, we consider a system where *N* individuals of two species are constrained to a one-dimensional space (a channel open at both ends) as shown in Fig 1B. We assume that the channel is always full and that the two species are segregated such that only one boundary separates the populations. Without loss of generality, we take *n* to describe the number of individuals belonging to the species on the left side of the lane (species 1). Contrary to the Moran model, the transition rates of the populations now depend on the spatial arrangement of the cells. A cell at any location can divide and produce a progeny, but death events only happen when a cell is pushed out of either end of the channel.

In this system, the relative species abundance delineates the location of the boundary that separates the species. When an individual cell grows, the boundary shifts right or left, as illustrated in Fig 1C. As *n* increases, the boundary between the two species moves to the right with probability *T* ^+^(*n* → *n* + 1); conversely, decreases in *n* result in the boundary moving to the left with probability *T* ^−^(*n* → *n* − 1). This competitive process continues until a sequence of jumps makes the boundary reach either end of the channel (i.e., *n* = 0 or *n* = *N*) with only one surviving species, which is said to fixate.

Like the Moran model, fitness differences modify the selection probabilities. The probability to select a specific individual *i* of species 1 for birth is 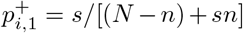, whereas the probability for individual *i* of species 2 is 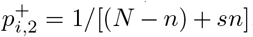. Following the birth, the progeny must make room for itself by pushing the cells on either side of its progenitor outwards. All cells in line are jointly pushed outwards, with the cell at the end of the channel getting removed from system by falling outside the channel.

In open-ended micro-channels, it has been experimentally observed that cells are more likely to grow in the direction of the closer channel opening because fewer cells need to be pushed in that direction. The probability for a cell to grow towards one of the openings was observed to scale linearly with the number of individuals between the cell location and the other opening [57]. Therefore, we assume that the probability that an individual cell at position *i* grows to the right is proportional to the number of individuals that are to the left of it *p*_left_ = (*i* − 1)*/*(*N* − 1) and vice versa *p*_right_ = (*N* − *i* − 1)*/*(*N* − 1). This spatial exclusion model is illustrated in Fig 1C.

Thus, the rate at which the boundary moves to the right in an interval of time Δ*t* is the sum of the rates for each individual of species 1 to grow to the right

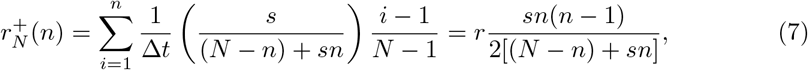

whereas the rate of the boundary moving to the left is

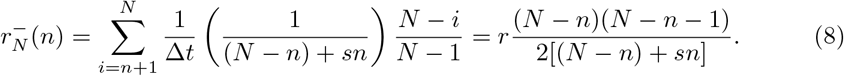

Here, we have rescaled the basal rate to *r* = 1*/*(*N* − 1)Δ*t*. These rates can be substituted in Eq 4-5 and solved to clarify the dynamics of the spatial exclusion model.

## Results

### Spatial exclusion gives rise to sharp sigmoidal fixation probabilities and exponentially fast MFPTs

We first consider a neutral case where species are functionally equivalent without fitness differences between them (*s* = 1) (e.g., two populations of cells whose phenotypic differences offer no upper hand or two identical lineages with a common ancestor). In contrast with the neutral Moran model, *A*(*x*) = [(*s* − 1)*x*^2^ + 2*x* − 1]*/*2[1 + *x*(*s* − 1)] and *B*(*x*) = [(*s* + 1)*x*^2^ − 2*x* + 1]*/*2[1 + *x*(*s* − 1)] for the spatial exclusion model. Although an analytical solution to Eq (4) is not available for *s* ≠ 1, it can be easily numerically integrated (see also S1 Appendix). Fig 2 shows the results of a comparison between the neutral Moran model and the spatial exclusion model.

**Fig 2.**
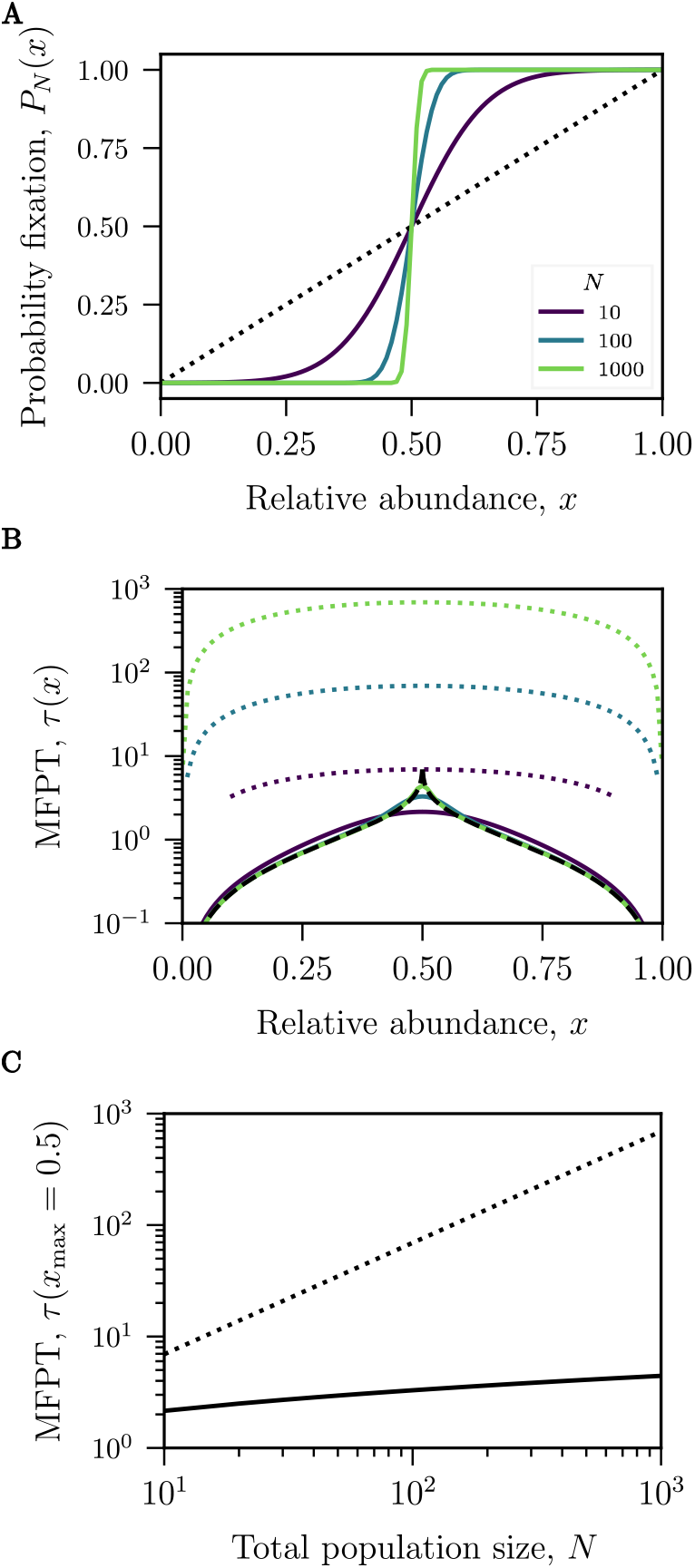
Competition outcomes for neutral dynamics. Fokker-Planck approximation for the fixation probabilities and mean-first passage times (MF-PTs) for the Moran model (dotted lines) and the spatial spatial exclusion model (solid lines) for various system sizes (*N* = 10, 100, 1000). (**A)** The fixation probability in the spatial model is sigmoidal around an inflection point at *x* = 1*/*2 and becomes increasingly steep, approaching a step-function at large *N*. The Moran model, in contrast, always predicts a linear fixation probability equal to the relative abundance. **(B)** For the spatial exclusion model, the MFPTs at varying *N* collapse onto each other for most relative abundances except around the inflection point. The dynamics away from *x* = 1*/*2 follow a deterministic path to fixation approximated by *τ*_det_ (dashed line). Notably, the MFPTs in the spatial model are significantly faster than those predicted by the Moran model. **(C)** The maximal MFPT of the Moran model is linear in *N* whereas the maximal MFPT of the spatial model grows substantially more slowly (sub-polynomial in *N*).

We find that the probability for a species to fixate in the spatial exclusion model as a function of its initial relative abundance is a sigmoidal function. This differs significantly from the linear dependence predicted by the Moran model (see Fig 2A). Any minority population (with a starting fraction *x <* 1*/*2), is much less likely to take over the population than in the Moran model. Conversely, any majority population (*x >* 1*/*2) is much more likely to succeed at fixating within the lane. The slope of this sigmoidal probability depends on the length of the microchannel (or the total population *N*) approaching a step function for large *N*. The inflection point is found at the equiprobable abundance *x*_max_ = 1*/*2, defined as the abundance at which both species are equally likely to take over the channel.

For the Moran model, the mean time to fixation grows linearly with the system size. For the spatial exclusion model, the mean time to fixation is exponentially shorter than that of the Moran model for all initial boundary positions (see Fig 2B). Interestingly, the MFPT curves for different *N* collapse onto each other away from the central peak at the equiprobable abundance of 1*/*2. This implies that the time to fixation, essentially, does not depend on system size unless the initial sizes of the two populations are closely balanced.

This independence of the dynamics on *N* can be heuristically explained by investigating the behaviour of Eq (3). For the spatial model, the first term on the right hand side (the drift term) is small, *A*(*x*) ≈ 0, around the peak of the MFPT. The dynamics around the peak are dominated by the second (diffusion) term on the right hand side of Eq (3), which scales like 1*/N*. Conversely, away from the peak, the drift term dominates the expression for large *N*. In this case, Eq (3) can be approximated by a first-order ODE

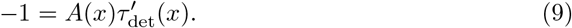

The solution to this equation

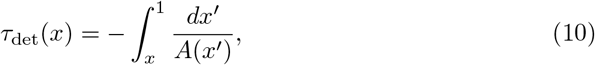

is the time to fixation for a process with a deterministic velocity *A*(*x*). For the neutral spatial exclusion model, using Eq 10, fixation occurs after time *τ*_det_(*x*) = − log |2*x* − 1|*/r*.

The displacement of the relative abundance is exponential in time when the change in relative abundance is governed by this deterministic velocity.

The mean-time to fixation of the spatial exclusion model substantially differs from the prediction of the Moran model, even close to the peak, where the MFPT depends on *N*. This difference between the two models for the MFPTs to fixation is most apparent at a starting fraction of *x* = 1*/*2 where *A*(*x*) = 0. In Fig 2C, the MFPT to fixation in the neutral spatial exclusion model grows only sub-polynomially with *N* rather than linearly as in the classical Moran model.

### Fitness differences break the symmetry of the fixation probability and engender longer MFPTs

So far, our description of the fixation times and probabilities has focused on neutral populations with similar or functionally identical species. However, more generally, phenotypically dissimilar species exhibit differences in their dynamics such as growth and death rates, efficiency of resource consumption, etc., which may impact their overall fitness in the environment. In this section, we investigate population fixation in the presence of a fitness difference, *s*, between two species. Although results are shown for *s >* 1, results for fitness differences *s*^*′*^ *<* 1 can be obtained by interchanging species 1 and 2 with *s* = 1*/s*^*′*^.

In the Moran model, the probability of fixation is heavily favored toward the species with a fitness advantage for even modest values of *s*, (Fig 3A)), with the probability of fixation being *p*(*x*) = (1 − *s*^−*xN*^)*/*(1 − *s*^−*N*^) for the fitter species [79]. Selection quickly skews the fixation probabilities so that the species with the advantage almost always fixates regardless of its initial abundance.

**Fig 3.**
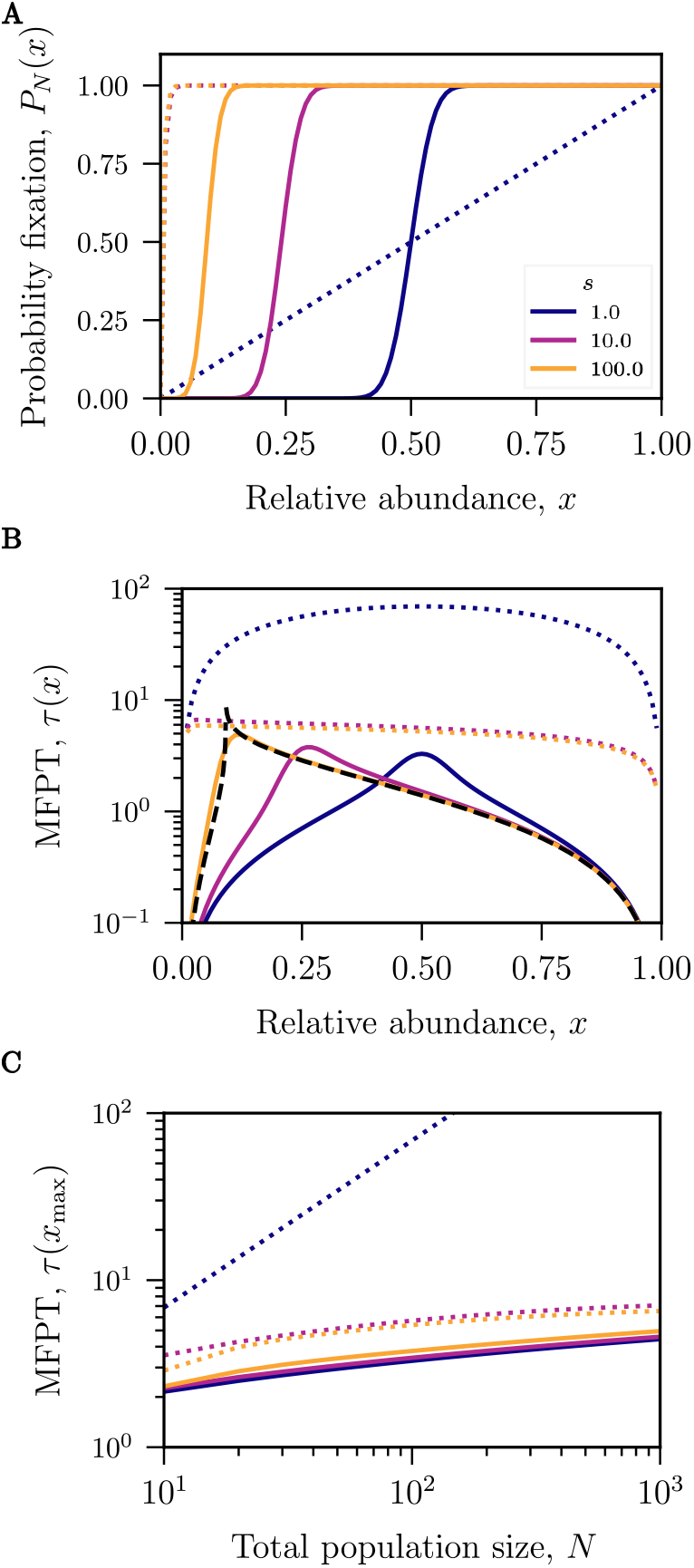
Competition outcomes for dynamics with fitness differences. Fokker-Planck approximation for the fixation probabilities and the mean-first passage times (MFPTs) for both the Moran model (dotted lines) and the spatial exclusion model (solid lines) with selection (*s* = 1, 10, 100). The relative abundance *x* is the abundance of the species with the selective advantage. **(A)** The fixation probabilities for both models with selection are shifted compared to neutral models, benefiting from a selective advantage. In plots (A) and (B) the system size is fixed at *N* = 100; however, the location of the inflection point does not differ for varying system size, only the steepness around the inflection point is affected as in the neutral models. In the upper left corner, the probability of fixation for the Moran model with *s* = 10 and *s* = 100 overlap. **(B)** In the spatial model, the maximum MFPT shifts with the inflection point and increases with fitness. In contrast, for the Moran model, the MFPTs tend to diminish with *s*. **(C)** The first passage time for the neutral (*s* = 1) Moran model exhibits linear growth in *N* ; however, this is the exception as the maximal first passage time for all other models grow sublinearly. These MFPTs all have very similar slopes irrespective of the fitness advantage. Still, they are shifted such that increased fitness leads to an increased time to fixation.

By contrast, similar fitness differences do not influence the population dynamics as markedly in the spatial exclusion model. Whereas the probability of fixation as a function of initial fraction *x* changes form drastically in the Moran model, from a linear function at *s* = 1 to a concave function without an inflection point for *s >* 1, within the spatial model, the shape of these curves remain relatively unchanged for all values of *s* (see Fig 3A). Rather, fitness differences shift the inflection point of the sigmoidal fixation probability curves towards lower initial relative abundances 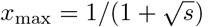, but a species is still significantly favoured to fixate when it’s relative abundance is above the inflection point. In other words, fitness differences in the spatial model offer a competitive advantage to the species with increased fitness, but this advantage is not as beneficial as the competitive advantage conveyed by a similar fitness difference in the Moran model. This is also reflected in the probability of fixation averaged over all initial abundances, see S1 Appendix, given that the average probability of fixation for the Moran model is higher than the average probability in the spatial exclusion model. Fitness advantages also impact the dynamics of the fixation. As shown in Fig 3B, higher fitness differences in the Moran model reduce the MFPT to fixation. Counter-intuitively, in the spatial exclusion model, although a greater fitness is generally associated with a higher likelihood of population fixation, increased fitness does not lead to faster fixations. Furthermore, the dependence of the MFPT on the relative fitness in the spatial model is location dependent and non-monotonic as shown in Fig 3B. For instance, the MFPT is maximal for populations initialized at abundances close to the equiprobable abundance 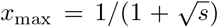, *τ* (*x*_max_) for any fitness (see Fig 3B), but this time increases with the increase in fitness difference between the species.

Accordingly, in the spatial exclusion model, the system’s longest timescale is determined by the competition dynamics around *x*_max_ where the probabilities of either species successfully taking over are roughly equivalent. The asymptotic scaling of this maximal MFPT with the system size *N* is similar to the asymptotic scaling in the neutral case (see Fig 3C). The non-linear slope of the MFPT on the log-log plot indicates that the growth of the MFPT is slower than a power law in *N* (this is further explored in discussions of Fig 5). The MFPT of the Moran model with selection also shows a sublinear growth in the population size, however, it appears in Fig 3C that this growth is slower than that of the spatial model.

Outside of this regime, there is a deterministic regime where the times are all almost identical. Other than the maximal MFPT, the times to fixation for various system sizes, *N*, collapse onto each other (see Fig 3C) in the deterministic regime. As in the neutral case, we can calculate a deterministic time

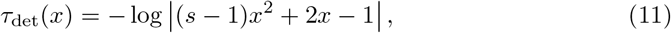

which is independent of system size and recovers the behaviour of the time of fixation (see Fig 3B). These species abundance dynamics correspond to a “tug-of-war”; the two species abundances fluctuate around the equiprobable abundance, with both species trying to take control of the the channel by their abundance. However, there is a quick switch to a deterministic takeover when one species becomes dominant enough at which point the growth in the dominant species abundance accelerates, causing the collapse in the abundance of the other species.

Given Eq 11, the asymptotic dependence of the MFPT on *s* far from the equiprobable abundance is logarithmic in 1*/*(*s* − 1). In other words, the MFPT away from the equiprobable abundance will asymptotically decrease with fitness differences. This is contrary to the behaviour of the MFPT at the equiprobable abundance, as it increases with fitness. Just as in the neutral case, the MFPT for most initial relative abundances is recovered using this deterministic approximation of the dynamics.

### Invasions are less likely to succeed due to spatial exclusion, but succeed on shorter timescales

As we have shown, the average time to fixation in a confined channel is sublinear in *N*. The fixated species remain dominant unless an external event, such as an invasion or mutation, perturbs the system introducing a new species variant that could compete against the established strain [80–82]. In microchannels, a mutation event at any location would introduce a new variant, but immigration is normally possible only at the edges. However, bacterial populations growing in wider microchannels have been shown to organize into parallel lanes aligning and growing along the axis of the channels. These aligned lanes permit rare immigration events from one lane into another previously fixated lane, which can be viewed as the invasion of a new species into a lane [57, 83].

We are interested in the probability and the mean time of a successful invasion wherein an individual cell of the invading species is inserted into the channel. Success of an invasion is defined as the invading species taking over and fixating within the lane/channel. In the well-mixed Moran model, the invasion of an individual from a new species into a previously fixated system results in an initial relative abundance of 1*/N* for the invading species. Derived in earlier sections, the probabilities of a successful invasion fixation for the Moran model (*P*_1_(1*/N*) = (1 + *s*^−1^)*/*(1 + *s*^−*N*^)) are depicted in Figs 2 and 3. The directional MFPTs of a successful invasion are solutions to Eq 5, shown in Fig 4C&D.

**Fig 4.**
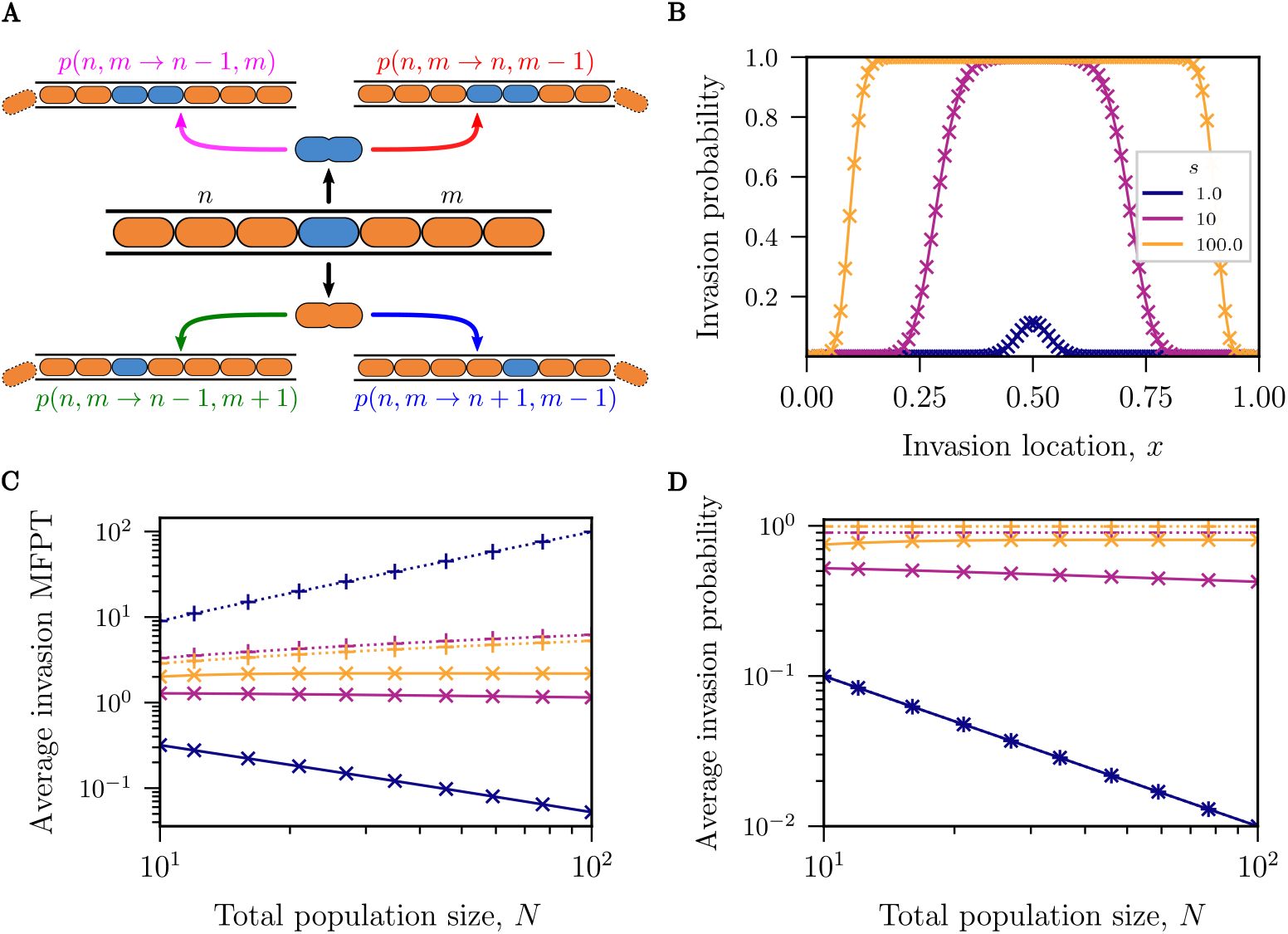
Invasion into a fixated channel. **(A)** Illustration of the invasion/mutation model wherein an invader (blue) with selective advantage *s* infiltrates a native population (orange). The invasion is shown at the center for illustrative purposes. (**B, C, D**): Numerical solutions to the Master equation for the spatial model (× connected with solid lines) and Moran model (+ connected with dotted lines). **(B)** The probability of successful invasion as a function of the insertion location (*N* = 100). Invasion events at the center of the channel are the most successful, but a selective advantage by the invading species can greatly increase the range of locales over which an invasion might succeed. **(C)** The average probability of successful invasion obtained by averaging the probability of successful invasion at all locations along the micro-channel. For *s* = 1, both models show similar behavior. **(D)** The average MFPT of a successful invasion in the spatial model decreases as population size increases, opposite the prediction of the Moran model. Moreover, as the selection advantage increases, so does the MFPT.

**Fig 5.**
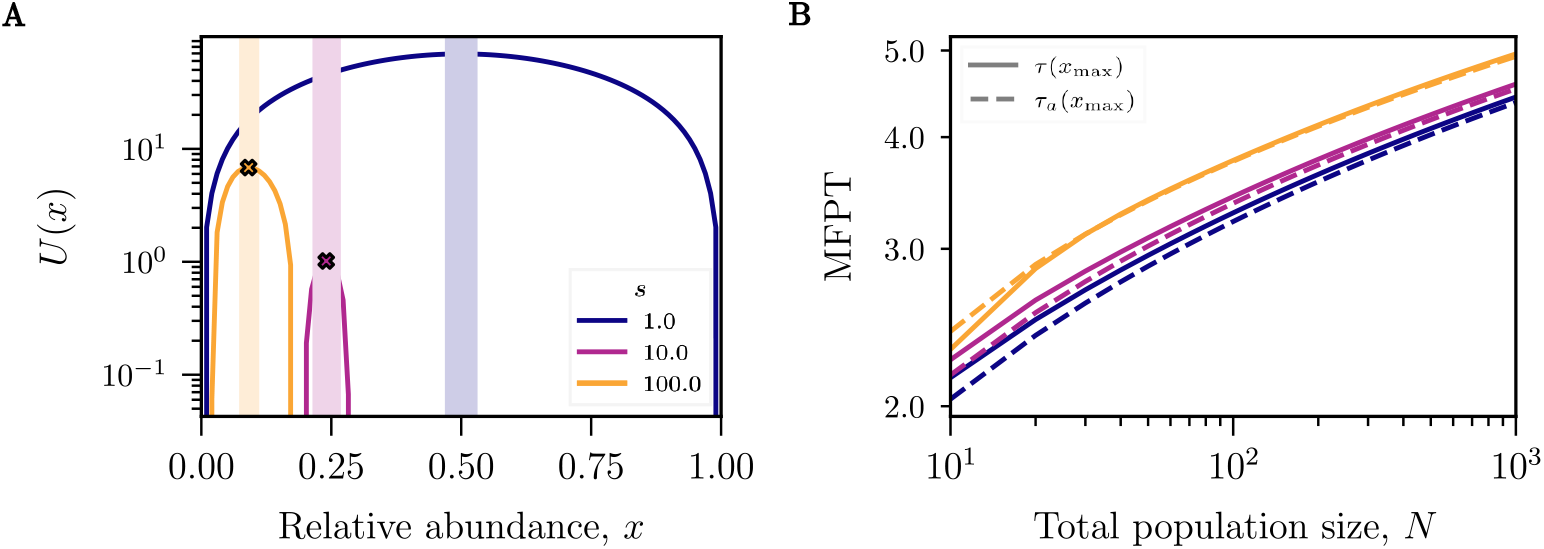
Heuristic approximation of the MFPT. **(A)** Graphs of the potential *U* (*x*) with the location of the maximum, 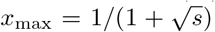, indicated at the peak of the curves. The shaded areas represent regions along the channel, *x*, where the stochastic behaviour heuristically dominates the dynamics (*N* = 100). **(C)** The maximal mean fixation time (solid lines) as a function of system size steadily increases for different fitnesses. The approximation *τ*_*a*_(*x*_max_) (dashed lines) is very close to the exact dynamics when a constant *τ*_dif_ ≈ 0.4 is added to the deterministic time *τ*_det_.

The outcome of an invasion in the spatial exclusion model, however, is strongly dependent on the initial location of the invasion event. Unlike the fixation problems studied in the previous section, the appearance of an immigrant in the one-lane model results in two rather than one inter-species boundaries as illustrated in Fig 4A. Nevertheless, the mathematical framework of competitive spatial exclusion described above allows us to model inter-species dynamics after such an invasion event, as shown in Fig 4A. The rules that determine the probability for a cell to be selected for birth and death are identical to those outlined previously. However, the system defined by the two inter-species boundaries is now two-dimensional and, accordingly, has added states with rates defining additional transitions. The expressions for these rates and the corresponding two variable master equation are outlined in S1 Appendix. Given that the state space for the spatial invasion model is now two-dimensional, we rely on numerical solutions for the discrete dynamics instead of solving the corresponding 2D Fokker-Planck equation.

We find that invasions happening about the center of the channel are the most likely to succeed in pushing out the established species, see Fig 4B. The probability that an invasion event succeeds increases with the fitness advantage of the invading species and successful invasions occur over a broader range of locations along the channel.

To provide a global measure of invasion success, we calculate the average invasion probability and the average MFPT to invasion, averaged over all possible initial locations of the invasion event. Surprisingly, this average invasion probability in the neutral spatial model is identical to the invasion probability of the neutral Moran model (*s* = 1) (i.e., it is equal to 1*/N* as shown in Fig 4C).

As demonstrated in Fig 4C, the probability of a successful invasion increases with fitness more sharply for the Moran model than for the spatial model. However, the probability of successful invasion remains low even for an invasive species with a ten-fold increase in fitness advantage, which is still less likely to fixate than the native species. This suggests that the spatial exclusion dynamics modeled here limit the competitive edge of strains with fitness advantages.

Although the probability for the invasive species to fixate is identical under neutral conditions for both the Moran model and the spatial exclusion model, the dynamics of fixation differs between the two as shown in Fig 4D. The MFPT for a successful invasion is larger for the Moran model than the spatial model in the neutral case. This is also true for for an invader with a fitness difference: successful invasions fixate more rapidly in the presence of spatial exclusion than in the well-mixed model.

Additionally, we find that the average MFPT of a successful invasion in the spatial model increases with fitness advantage, reminiscent of the behaviour of the MFPT Fig 3B&C. This is contrary to the intuition that higher selective advantage should accelerate the fixation of the invasive species because the fitter invader grows more rapidly. A successful invasion must go from the two interspecies boundaries to one interspecies boundary before taking over the channel; thus, the MFPT is limited by the one boundary MFPT which increases with fitness advantage as shown in Fig 3B&C.

### Quantitative investigation of different dynamical regimes in large systems competing in a tug-of-war

The dependence of the fixation time on system size *N* is of importance when interpretating experimental results that commonly probe only the transient composition of evolving ecosystems. In Fig 3C we have numerically seen that the asymptotic behaviour of the MFPT to fixation shows subpolynomial growth in *N* for the spatial exclusion model. In this section, we investigate the large *N* scaling of MFPTs - a regime relevant for many experimental systems that are often comprised of large numbers of individuals.

The one-dimensional Fokker-Planck equation for the MFPT to fixation, Eq 3, may be rewritten as

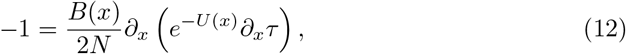

where

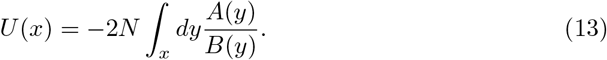

*U* (*x*), sometimes referred to as the Fokker-Planck potential, describes an effective potential landscape in which the boundary between the two species moves. A general compact integral form of the solution to Eq 3, which is shown in S1 Appendix, can be evaluated for different potentials representing different population dynamics.

For the model defined in Eq 7 and Eq 8, *U* (*x*) is a unimodal distribution with an unstable maximum found at 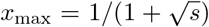 for *s* ≥ 1 (see Fig 5A). Thus, unlike the more familiar problem of calculating the MFPT to cross over a potential barrier (Kramer’s Theory) [75], here we are interested in finding the MFPT to descend from a potential peak starting at an unstable point. This subtle distinction means that the saddle-point approximation - which is commonly employed for getting the scaling of the MFPT in Eq 3 - is inadequate in this case.

To derive an approximation for the asymptotic behaviour of the fixation MFPT, we heuristically separate the space into regions of predominantly deterministic or stochastic dynamics. As shown in S1 Appendix, the boundaries between these two regions naturally emerge from the integral form of the MFPT as the solutions to the equation

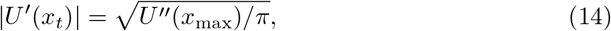

and are depicted in Fig 5A. The boundaries 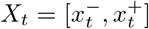 contain the region around *x*_max_ where the dynamics are dominated by the stochastic diffusion with the location-dependent diffusion coefficient *B*(*x*) arising from a ‘tug-of-war’ dynamics between the two species. Conversely, outside of this region, the dynamics are well approximated as deterministic and are dominated by an ecological drift with a location-dependent velocity *A*(*x*).

Thus, if the initial relative abundance of species 1 lies in the deterministic region *x* ∉ *X*_*t*_, the time to fixation is well approximated by the deterministic time *τ*_det_(*x*) of Eq 10. On the other hand, if the initial relative abundance is in the stochastic region *x* ∈ *X*_*t*_, its motion is at first dominated by diffusion until it reaches one of the boundaries within a time 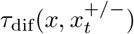, after which it can continue along a deterministic trajectory.

The approximate mean time to fixation from within the stochastic region can be written as

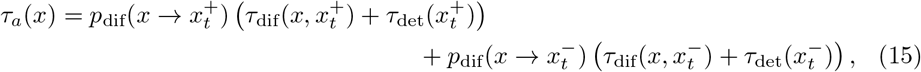

where 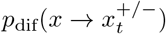 is the probability that the relative abundance *x* diffuses to one of the boundaries at 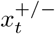.

For illustrative purposes, we examine the fixation time starting at *x* = *x*_max_ given that there is an equal probability to diffuse to either bound from this position. For the neutral spatial exclusion model, the maximal MFPT is

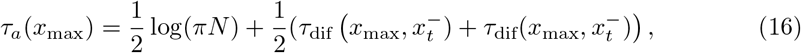

which is shown in Fig 5B. As shown in S1 Appendix), numerically *τ*_dif_ ≈ 0.4 and is independent of *N* or other parameters of the model (further analysis is needed to determine the constant analytically). The spatial exclusion model with selection also exhibits a 𝒪 (log *N*) scaling. For all selective advantages, we find that the difference between the deterministic solution and the full solution is, somewhat surprisingly, a constant (see S1 Appendix).

## Discussion

Spatial exclusion significantly alters the competitive dynamics between species in densely populated bacterial communities. Here, we have studied the competition between two species of bacteria confined to an open 1D microchannel as a starting point to understanding bacterial competition within more complex confined geometries. This required the development of a spatial exclusion model that explicitly accounts for the mechanical exclusion between cells, in contrast to non-spatial well mixed models such as the paradigmatic Moran model.

We find that the probability of species fixation in the spatial exclusion model shows a much sharper sigmoidal dependence on the initial relative abundance in contrast to the Moran model where the corresponding probability is equal to its initial relative abundance. The inflection point of the sigmoidal curve - where the probabilities of fixation of either species are equal - is located at the initial abundance *x* = 1*/*2 for neutral populations without selective advantage (*s* = 1), but shifts to lower initial abundances of the fitter species 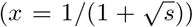 at higher values of selective advantage. With an increase in the population size, *N*, the sigmoidal curve approaches a step function, effectively setting a threshold in the initial abundance above (below) which the cells will always (never) fixate.

The mean fixation times in the spatial exclusion model are sped up in comparison to the predictions of the Moran model by up to several orders of magnitude. Interestingly, these fixation times have a very weak dependence on the total population size for most initial abundances, and are dominated by a quasi-deterministic exponential escape towards fixation. Only for the initial abundances near the inflection point of the fixation probability curve, where the maximum timescales are observed, does the fixation time show a weak dependence on population size that is well approximated by an asymptotic logarithmic scaling in population size.

Although the species with a selective advantage maintains a competitive advantage within our spatial exclusion model, this competitive advantage does not affect the fixation dynamics to the same extent as it does within the Moran model. Indeed, within the spatial exclusion model, fixation occurs at lower initial relative abundances of the fitter species than in the neutral model. However, the competitive advantage a bacterial species gains by being more fit than its competitor is significantly less in this constrained environment than in a well mixed system devoid of spatial limitations. Although births of the more fit species happen more frequently, the spatial organization of the cells make deaths of the more fit species happen more often as well (as they fall out of the channel) resulting in a reduction in the competitive advantage compared to the well-mixed model. Mechanisms that convey selective advantages can be difficult to maintain due to their strain on metabolism and increased resource costs [84, 85]. Furthermore, competition between species with higher fitness differences show longer fixation times despite the fact that one of the species has a greater likelihood of dominating the channel. The slower fixations are owing to slower diffusion relative to drift around the point of equiprobable abundance. This means that not only must a species preserve a potentially more resource intensive strategy to gain an advantage, but it must sustain this strategy for longer to insure it dominates. Thus, the overall competitive advantage provided by an increase in fitness must be weighed against these two competing effects.

Once one species fixates, competition is halted until another species is reintegrated into the system through an invasion process. Overall, an invader with a selective advantage increases the likelihood that an invasion at any location is successful, with the invasions most likely to succeed occurring in the middle of the channel. Similar to the Moran model, the probability of a successful invasion (averaged over all initial conditions in the channel) decreases as the total population size increases. However, contrary to the Moran model, the mean invasion times averaged over the initial channel location decrease with system size. These fixation times accelerate in longer channel because successful invasions with larger population sizes are reliant on the the invader species remaining near the center of the channel, which fixates more rapidly. Consequently, successful invasions in longer channels happen less frequently, but those that are successful are more rapid on average.

In recent years, microfluidic monolayer devices (MMD) - such as mother machines, chemostats, etc. - have been designed to study single cell bacterial growth and generational dynamics [66, 86, 87]. Our results model the behaviour of competing populations in a single-lane, open chemostat and could be directly tested within such a device. However, our spatial model can also be extended to more complex 2D MMDs that support multi-lane channels. Pill-shaped bacteria, such as *E. coli*, are observed to grow constrained to 1D lanes within wide, open-ended microchannels [57, 58, 88]. As a first approximation, the larger channels can be viewed as many 1D lanes that interact through rare immigration events. In this simplified view, the dynamics of channel fixation can be decoupled from the lane fixation if the time between lane invasions is longer than the fixation time within a lane. Accordingly, future work will combine the probability of a successful invasion found in Fig 4 with the rate of invasion to model the 2D competition as in *Koldaeva et al*. [57].

For the 1D channel with fitness differences, we showed that one species is almost deterministically favoured to out-compete the other when the initial relative abundances are not close to the equiprobable abundance. By contrast, for initial abundances in a region close to the equiprobable abundance, the dynamics is dominated by stochastic diffusion. Either species is, roughly, equally likely to take control of the channel within this region with the two opposing species competing in a tug-of-war. Under the heuristic assumptions concerning the regions of dominant dynamics, we find that the tug-of-war concludes with one species outcompeting its rival in times that scale at most in 𝒪 (log *N*) when starting in the diffusive regime. This is significantly different from the paradigmatic Moran model which predicts a 𝒪 (*N*) fixation.

More generally, we have developed a heuristic framework to approximate the asympotic dependence (at large *N*) of the mean first-passage times in a tug-of-war process. However, a (more) exact analytical solution to the Fokker-Planck equations of these dynamics may be possible. For instance, various asymptotic expansions utilizing Watson’s Lemma may provide satisfactory limits for calculating the MFPT integrals [89]. Another promising future direction is a description of the bacterial spatial exclusion as a single-file dynamics model. Single-file dynamics are concerned with the motion of many particles along a line and can describe the mean squared displacement (MSD) of particles in a channel [90], which is often slow. However, translating our spatial model to an appropriate single-file dynamics model may uncover faster displacements akin to our predictions.

The heuristic framework for calculating the asymptotic MFPTs is of general interest and has applications well beyond bacterial population dynamics. The counter-intuitive results concerning the competition of the species are due to these tug-of-war dynamics, which may be applied to other systems with drift and diffusion terms leading to a concave effective potential as in Fig 5A. Our analysis may be applied to other systems whose dynamics are equivalent to the diffusion of a particle descending an effective potential. For instance, the transport of organelles and other cellular cargo has been described by a tug-of-war wherein competing sets of molecular motors pull in opposite directions with the drift depending on the number of motors on either side of the cargo, much like our model [91]. Moreover, competition between populations of cancer and healthy cells display a tug-of-war effective potential that recovers a probability of cancer development that is sinusoidal as a function of the initial relative abundance of the cancerous cells, like in the spatial exclusion model [92]. Our methodology can predict the probabilities of and mean-time to a clinical outcome to determine the rapidity of the disease.

In summary, we have shown that explicitly incorporating spatial interactions arising from cell growth and division within dense bacterial populations can have important consequences for both the overall composition and the rate of species exclusion from the system. Our results provide insights into the processes involved in the formation and maintenance of complex bacterial ecosystems such as biofilms, intestinal flora, or various persistent infections. Likewise, the techniques developed here may more broadly be applied to a range of competitive dynamical systems from cellular transport to cancer.

## Supporting information

### S1 Appendix

#### Derivations of the probabilities and MFPT to fixation

We review the calculation of the probability and exact mean first-passage time for the discrete models discussed in this main text and the derivation of their corresponding continuous Fokker-Planck approximations. The averaged quantities and invasion dynamics discussed above are detailed and discussed.

## Acknowledgments

T. Ma and J. N. Milstein acknowledge funding from an NSERC Discovery Grant and a New Frontiers in Research Fund Exploration Grant. A. Zilman acknowledges support from an NSERC Discovery Grant. The authors are indebted to the members of the Milstein, Zilman and Goyal labs for numerous discussions.

